# A tetracycline antibiotic minocycline prevents early aging phenotypes in mice heterozygous for RP58

**DOI:** 10.1101/2021.09.18.460879

**Authors:** Tomoko Tanaka, Shinobu Hirai, Hiroyuki Manabe, Kentaro Endo, Yasumasa Nishito, Hiroko Shimbo, Hikari Yoshitane, Haruo Okado

**Affiliations:** Department of Psychiatry & Behavioral Science, Tokyo Metropolitan Institute of Medical Science, Tokyo, 156-8506, Japan; Department of Basic Medical Science, Tokyo Metropolitan Institute of Medical Science, Tokyo, 156-8506, Japan; Laboratory of Neural Information, Graduate School of Brain Science, Doshisha University, Kyoto, 610-0394, Japan; Center for Basic Technology Research, Tokyo Metropolitan Institute of Medical Science, Tokyo, 156-8506, Japan

**Keywords:** aging, DNA damage, DNA repair, inflammation, RP58

## Abstract

**Summary:** In humans, cognitive and motor functions develop in association with maturation, followed by a decline in advancing age. In this study, we aimed to provide a method for extending healthy lifespan by preventing age-related phenomena. Therefore, we focused on RP58, a transcriptional repressor, whose expression is reduced during aging in the human cortex. In the *Rp58* hetero-knockout (KO) mice, object location memory was impaired even at 4–5 months, while it was normal in the wild-type mice at 4–5 months but was impaired at 12–18 months. These results indicate an early onset of impaired spatial memory in the mutant mice. As the underlying mechanism, the *Rp58* hetero-KO mice showed early onset of DNA damage accumulation and microglial activation in the dentate gyrus due to a DNA repair defect that is generally observed with aging. As another hallmark of aging, we focused on mitochondrial function and detected mitochondrial abnormalities in the *Rp58* hetero-KO mice at 4–5 months. Notably, continuous treatments with minocycline, a neuroprotective and anti-inflammatory antibiotic, prevented the facilitation of age-related phenomena in the *Rp58* hetero-KO mice. These results suggest the availability of the *Rp58* hetero-KO mice as a novel mouse model of human-like early aging and provide a therapeutic strategy to prevent age-related phenomena using minocycline.

**Highlight:**

- *Rp58* hetero-KO mice exhibit early aging phenotypes including impairment of spatial cognition
- *Rp58* hetero-KO mice show early accumulation of DNA damage due to a defect in the DNA repair system
- Treatment with minocycline prevented cognitive dysfunction observed in *Rp58* hetero-KO mice

## Introduction

Aging increases the risk for neurodegenerative disorders, cancer, and other chronic disorders in humans. Medical technology is advancing rapidly, and life expectancy is increasing; therefore, it is imperative to develop strategies to prevent the onset of disease and/or to maintain the quality of life after the onset of disease. Symptoms such as an age-related decline in physiological functions are observed throughout the body. These symptoms are associated with each other and worsen with age [1]. Thus, age-related symptoms not only progress in an age-dependent manner but are also further affected by indirect effects from other age-related phenomena, *e*.*g*., cytokines as senescence-associated secretory phenotypes and insomnia-induced systemic function decline. It is therefore difficult to distinguish between cause and effect and to identify target mechanisms for therapy. Recently, to better understand the aging process, mouse models of human aging have been utilized to examine every aspect of the observed symptoms separately [2]. For example, mutations in the human mitochondrial *polymerase gamma* (*POLG*) gene have been frequently found in patients with mitochondrial disease, Parkinson’s disease, and cancer. Mice with mutations in the *Polg* gene exhibited premature aging phenotypes, such as mitochondrial dysfunction and hyperinflammatory innate immune status [3]. Furthermore, the *amyloid precursor protein* (*App*) knock-in mice with Swedish, Iberian, and Arctic mutations (*App*^*NL-G-F/NL-G-F*^ mouse), known as the Alzheimer’s disease model, exhibited Aβ_42_-dominant plaque deposition concomitantly with neuroinflammation and memory impairments [4]. These phenotypes are observed in some dementia patients but not in others. Therefore, it is imperative to generate a new mouse model that includes other aspects of age-related symptoms to propose new anti-aging therapies. Moreover, we should emphasize that therapeutic strategies for the prevention of age-related symptoms would be more efficient than those for the recovery from severe conditions of such symptoms.

It has been reported that the expression of a series of genes is reduced in the aged human cortex [5]. In order, from top to bottom according to the age-related decline in expression level, these are SCN2B, a voltage-gated Na^+^ channel; MAP1B, a microtubule-associated protein; and RP58, a transcriptional factor. RP58 is a 58-kDa transcriptional repressor protein also known as Zinc finger and BTB domain-containing protein 18 (ZBTB18) [6]. We previously generated *Rp58* knockout (KO) mice and demonstrated that RP58 plays a critical role in the formation of the cerebral cortex [7-9]. RP58 is highly expressed in the neonatal brain but its expression continues after the brain maturation [10]. RP58 represses the expression of Id4 [11], which promotes neural stem cell quiescence in the adult dentate gyrus [12]. Also in human, some cases of developmental disorders with *RP58* mutations have been reported [13], while only a few studies on the role of RP58 after adolescence have been conducted. It would be useful to determine the impact of a decrease in RP58 expression on the development and progression of age-related symptoms.

Here, we clarified that the *Rp58* hetero-KO mice show early DNA damage accumulation due to a DNA repair defect that is generally observed with aging. Inflammation was observed concurrently with the accumulation of DNA damage in the *Rp58* hetero-KO mice. These phenomena may result in early biological aging including impairment of spatial cognition. Moreover, we also evaluated the efficacy of minocycline, which has neuroprotective and anti-inflammatory effects [14, 15], against the aforementioned phenomena. Our research suggests that the *Rp58* hetero-KO mice can be used as a novel mouse model of human aging, and it also provides a potential therapeutic strategy to prevent age-related phenomena by using minocycline.

## Results

### Hetero-KO of Rp58 induces early impairment of cognitive function during aging

To extend the healthy lifespan by preventing age-related phenomena, we aimed to generate a novel mouse model of human aging and to search for a therapeutic strategy to prevent the age-related phenomena observed in the model. Given that expression levels of *Rp58* have been reported to be decreased in the aged human cortex [5], we focused on *Rp58* hetero-KO mice as a potential mouse model of human-like early aging. We first investigated if the mutant mice exhibit an early onset of impaired spatial memory, which is an age-related phenomenon. Mice at 2, 4–5, and 12–18 months of age were respectively used to represent adolescence (young), adulthood, and old age in the object location test (Figure 1A) [16]. Young wild-type mice preferred a familiar object at a novel place to one at a familiar place, as evidenced by a discrimination index with significant difference from zero. A discrimination index in the wild-type mice is close to zero at 12–18 months but not at 4–5 months (Figure 1B). In contrast, object location memory in the *Rp58* hetero-KO mice was impaired even at 4–5 months, as evidenced by a discrimination index close to zero. These results indicate that reduced expression of *Rp58* accelerates aging and influences cognitive function. To determine an impaired process of spatial memory in the *Rp58* hetero-KO mice, we recorded the theta power in the hippocampal CA1 region, because the hippocampal CA1 region is crucial for spatial learning and the theta power selectively reflects the successful encoding of new information (Figure 1C). In the wild-type mice at 4–5 months, the average level of theta power was increased during the 2 seconds before and after the time of contact with objects on day 2, *i*.*e*., when the objects were introduced for the first time. However, this induction in the theta power was significantly attenuated on day 2 in the *Rp58* hetero-KO mice at 4–5 months. The decreased theta power in the *Rp58* hetero-KO mice was observed not only on day 3 but also on day 2, indicating that the *Rp58* hetero-KO mice had a defect in learning the place of the object rather than in recalling the memory.

**Fig. 1.**
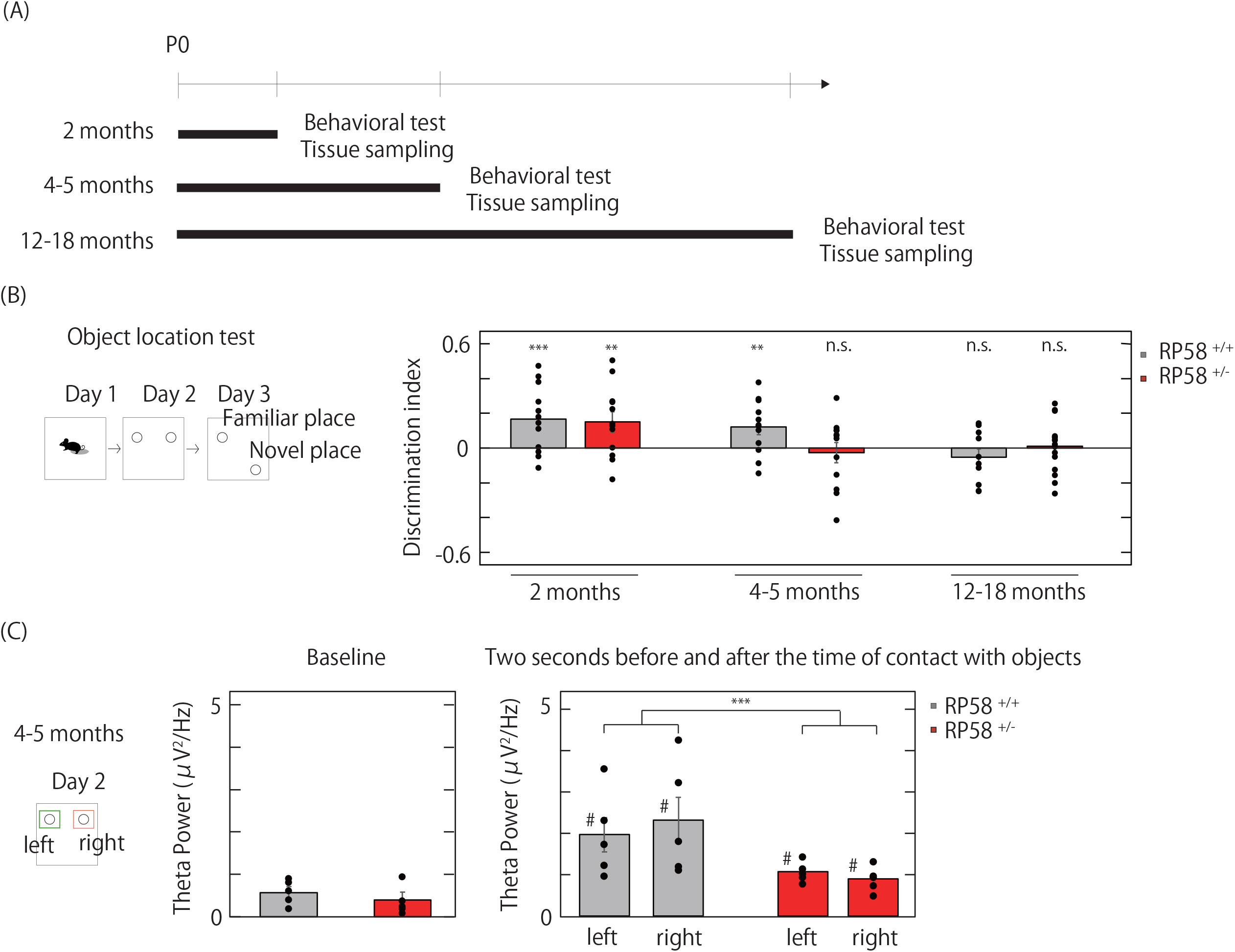
Spatial memory and hippocampal function in the *Rp58* hetero-KO mice (A) Experimental schedule. (B) Spatial memory test of the wild-type and the *Rp58* hetero-KO mice during aging. Data from the object location test were calculated and converted to the discrimination index (see Materials for details). *p < 0.05, **p < 0.01, ***p < 0.001, One sample t-test vs zero in the discrimination index; n = 12 for both genotypes at 2 or 4-5 months, n = 10 for wild-type mice at 12–18 months, and n = 16 for the *Rp58* hetero-KO mice at 12–18 months. (C) Theta power in the hippocampal CA1 region of the wild-type and the *Rp58* hetero-KO mice at 4–5 months on day 2 of the object location test. ^#^p < 0.05, Paired t-test vs baselines for each genotype, n = 5 for each group. ***p < 0.001, Paired t-test; n = 10 (two objects per mouse and five mice per group) for each genotype.

### Hetero-KO of Rp58 induces early accumulation of DNA damage in the mossy cells of the dentate gyrus

To identify which types of cells in the hippocampus contribute to the effect of the *Rp58* hetero-KO on the spatial memory, we performed immunohistochemistry of the adult dentate gyrus by using an anti-RP58 antibody [17]. We found that RP58 protein was highly expressed in NeuN-positive cells (neurons) of the hilus region (most probably the mossy cells) and also in NeuN-positive cells of the granule cell layer (Figure 2A). The mossy cells play a crucial role in spatial memory by receiving convergent excitation from the granule cell and returning bilateral, widespread, and divergent excitation to the granule cell [18, 19]. The localized expression of RP58 protein in the mossy cells led us to examine the possibility that early aging onset was observed in the mossy cells of the *Rp58* hetero-KO mice. Among the multiple symptoms of senility, we examined the accumulation of DNA damage, a contributor to the progression of aging, by detecting ssDNA in the mossy cells, because ssDNA is generated after DNA damage. In the wild-type mice, the number of ssDNA-positive neurons gradually increased from 2 months to 12–18 months (Figure 2B). In the *Rp58* hetero-KO mice, apparent ssDNA accumulation was detected even at 4–5 months, while no significant effects were observed at 2 months and at 12–18 months, indicating that the *Rp58* hetero-KO mice showed early onset of DNA damage accumulation in the mossy cells. The early accumulation of DNA damage in the mutant mice was also confirmed by another DNA damage marker—gamma-H2AX, a phosphorylated form of H2AX protein at Ser139 [20] (Figure 2B). Therefore, *Rp58* likely has a key role in the prevention of DNA damage accumulation in the mossy cells of the dentate gyrus of the adult hippocampus during the aging process.

**Fig. 2.**
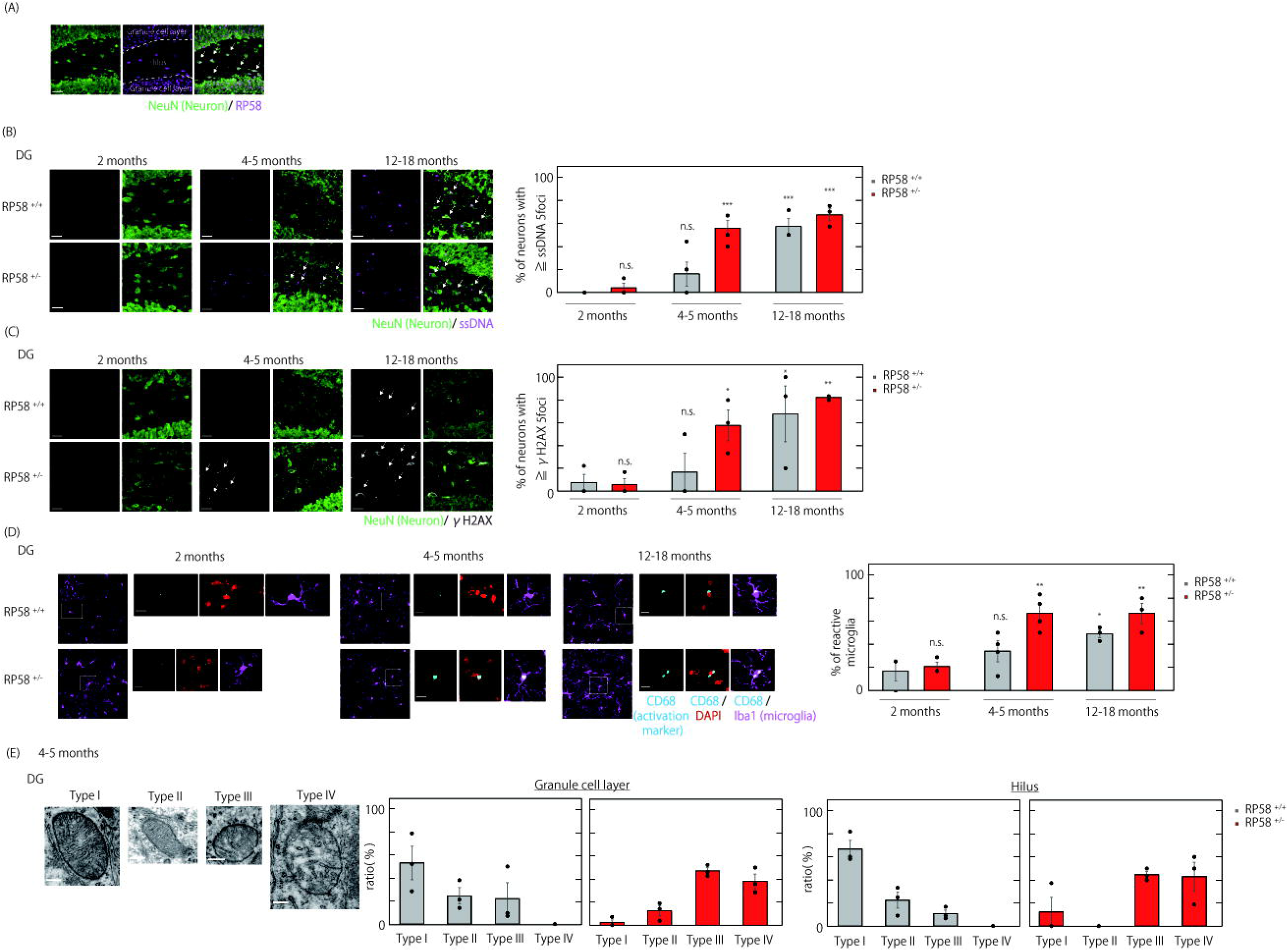
Cellular mechanisms of DNA damage in the *Rp58* hetero-KO mice (A) Localization of RP58 protein in the dentate gyrus in adult mice. (B) Accumulation of ssDNA (indicated by arrows) as a DNA damage marker in neurons in the dentate gyrus of the wild-type and the *Rp58* hetero-KO mice.. ***p < 0.001, Dunnett’s test vs wild-type mice at 2 months; n = 3 for both genotypes at 2 or 12–18 months, and n = 4 for both genotypes at 4-5 months. (C) Accumulation of gamma-H2AX protein (indicated by arrows) as a DNA damage marker in neurons in the dentate gyrus of the wild-type and the *Rp58* hetero-KO mice. *p < 0.05, **p < 0.01, Dunnett’s test vs the wild-type mice at 2 months; n = 3 for each group. (D) Microglial activation detected by labeling microglia with Iba1 (magenta) and activation with CD68 (cyan) in the dentate gyrus of the wild-type and the *Rp58* hetero-KO mice. *p < 0.05, **p < 0.01, Dunnett’s test vs the wild-type mice at 2 months; n = 3 for both genotypes at 2 or 12–18 months, and n = 4 for both genotypes at 4-5 months. (E) Morphological abnormalities of mitochondra of neuronal origin in the dentate gyrus of the wild-type and the *Rp58* hetero-KO mice. Based on their ultrastructural appearance, detected mitochondria were classified into one of four categories as was classified in the previous study [37]. Granule cell layer: chi-square = 284.837, degrees of freedom (df) = 3, p < 0.001; hilus: chi-square = 359.525, df = 3, p < 0.001, Pearson’s chi-squared test; n = 3 for each group.

### Activation of microglia and chronic inflammation in hetero-KO of Rp58

We subsequently focused on the microglia in the hilus region of the dentate gyrus because the accumulation of DNA damage in neurons is associated with abnormal activation of microglia in the periphery of the damaged neurons [21]. Microglia show an exaggerated response to the secondary inflammatory stimulus than the first one, and these extremely responsive microglia, called primed microglia [22, 23], become increasingly common with age [24]. For example, the primed microglia were found to be accumulated in a DNA repair-deficient mouse model which displays accelerated aging in multiple tissues including the central nervous system [25]. In the present study, we examined the onset of abnormal activation (also known as “priming”) of microglia in the hilus of the *Rp58* hetero-KO mice. In the wild-type mice, abnormally activated microglia cells that were stained not only by Iba1, a microglia marker, but also by CD68, an activation marker, gradually increased from 2 months to 12–18 months (Figure 2D). In the *Rp58* hetero-KO mice, abnormal accumulation of activated microglia was observed even at 4–5 months, while no significant effects were observed at 2 months and at 12–18 months, indicating that the *Rp58* hetero-KO mice exhibited another early aging phenotype.

The early onset of microglia activation in the *Rp58* hetero-KO mice raised the possibility that chronic inflammation in the brain was induced by mutation even in the early life stage. To test this possibility, we examined gene expression profiles in the hippocampus of the *Rp58* hetero-KO and the control wild-type mice in adulthood (4–5 months) and old age (12–18 months). Among the genes whose expression level was down-regulated in the aged wild-type mice, 341 genes were down-regulated in the adulthood *Rp58* hetero-KO mice when both were compared with the adulthood wild-type mice. Conversely, the expression levels of 103 genes were up-regulated both in the adulthood *Rp58* hetero-KO and the aged wild-type mice (Supplemental figure 1A). In the pathway enrichment analysis, these 103 up-regulated genes included *Cxcl10, Oas2, Oas1a*, and *Oasl2* (Supplemental figure 1B), which are associated with a pathway of immune response interferon gamma action on extracellular matrix and cell differentiation (Supplemental figure 1C).

### Ultrastructural changes in the mitochondria of Rp58 hetero-KO mice

To ensure the availability of the *Rp58* hetero-KO mice as a mouse model of human aging, we focused on the function of mitochondria. Mitochondria have a crucial role in antigen presentation and innate immune reactions and serve as “immunometabolic hubs” [26, 27]. The abnormalities in mitochondria have been shown to be age-related [28]. In this study, we categorized the mitochondrial structure into four types, Type I (normal) to Type IV (abnormal ultrastructure), as explained in the Methods section. In the *Rp58* hetero-KO mice at 4–5 months, most mitochondria in neurons of the granule cell layer and the hilus region were categorized as Type III and IV, while in the wild-type mice, most were categorized as Type I to III (Figure 2E). It should be emphasized that an exclusively abnormal type of mitochondria categorized as Type IV, which showed i) swollen and deficient cristae or ii) swollen cristae and a discontinuous outer membrane, were detected only in the *Rp58* hetero-KO mice (Figure 2E).

Collectively, the *Rp58* hetero-KO mice showed early onset of several aging phenotypes, such as impairment of spatial memory, DNA damage accumulation, microglial activation, and abnormal mitochondrial ultrastructure in the dentate gyrus. We propose that the newly generated aging mouse model can mimic RP58 decline and cognitive dysfunction in humans.

### Hetero-KO of Rp58 impairs the process of DNA repair

We aimed to propose an effective therapeutic strategy for human aging by exploring a therapeutic approach for our new aging mouse model. Based on a predicted molecular mechanism of abnormalities in the *Rp58* hetero-KO mice, we focused on the process of DNA repair. DNA is continually damaged by exogenous and endogenous sources, but these damages are not serious in the normal DNA repair state. On the other hand, impairment of the DNA repair pathway leads to accumulation of DNA damage, which has a harmful effect on adult health and the onset of pathology [29, 30]. Given that the *Rp58* hetero-KO mice show the early accumulation DNA damage,, we explored whether the *Rp58* hetero-KO mice exhibit any defects in the DNA repair pathway by monitoring recovery from irradiation-induced DNA damage. ssDNA-positive neurons were detected in the granule cell layer of the dentate gyrus 1 hour after irradiation in the wild-type mice at 2 months when they were exposed to X-ray (0.3 Gy) (Figure 3B) but not in the wild-type mice without irradiation (Figure 2B). The number of ssDNA-positive neurons gradually decreased from 1 hour to 24 hours after irradiation (Figure 3B). In the *Rp58* hetero-KO mice, a similar extent of ssDNA accumulation was observed 1 hour after irradiation; however, ssDNA accumulation was still observed at 24 hours after irradiation. The residual DNA damage even at 24 hours after irradiation was corroborated by another DNA damage marker, gamma-H2AX, in the *Rp58* hetero-KO mice (Figure 3C). Also, the ratio of abnormally activated microglia cells in the granule cell layer remained unchanged, even at 24 hours after irradiation (Figure 3D). These data strongly support our idea that RP58 plays a pivotal role in DNA repair. We thus demonstrate that the *Rp58* hetero-KO mice show early accumulation of DNA damage during aging (Figure 2) due to a defect in the DNA repair system (Figure 3); therefore, they exhibit early aging phenotypes including impairment of spatial cognition (Figure 1).

**Fig. 3.**
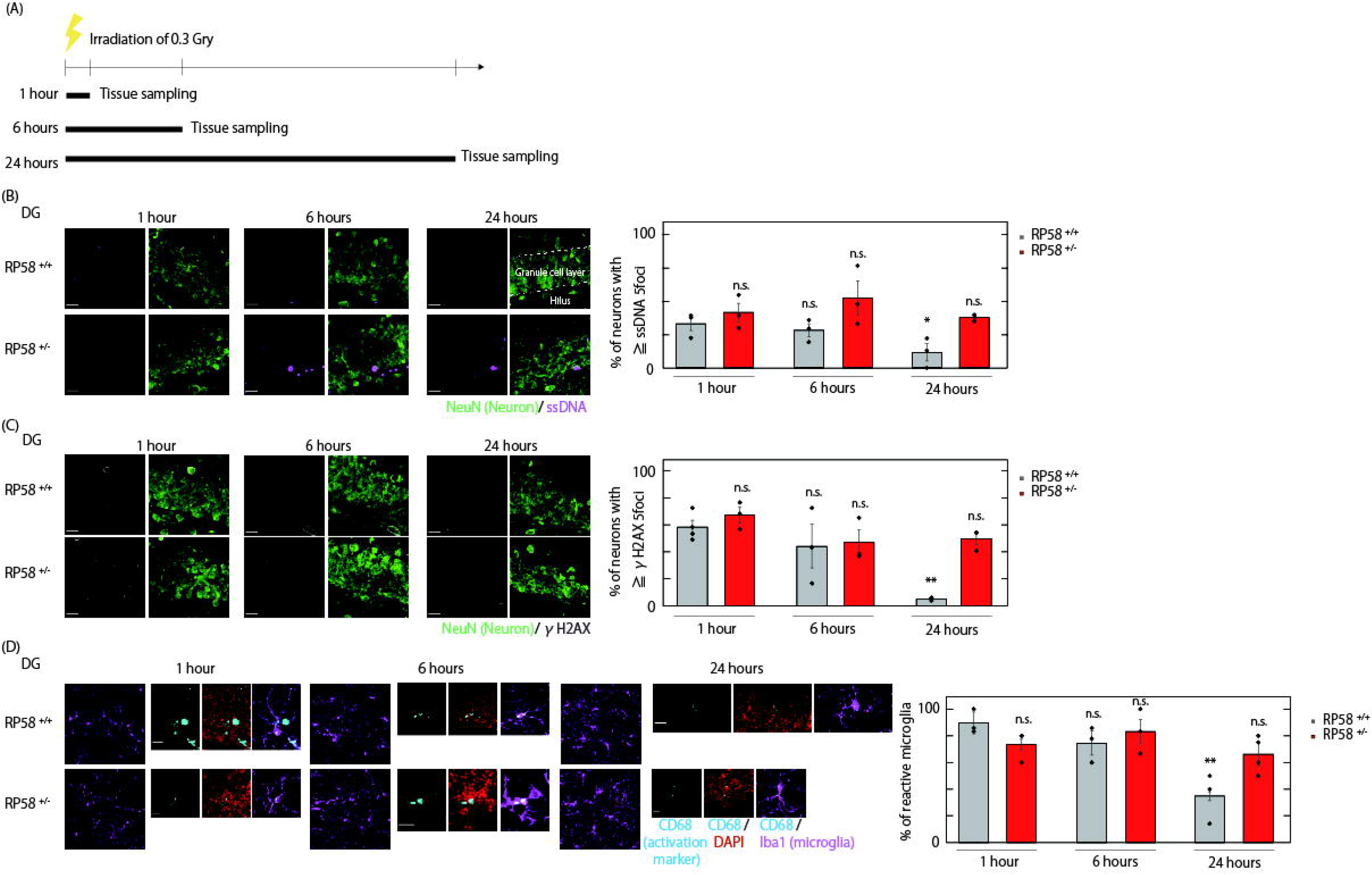
Dysfunction of DNA repair in the *Rp58* hetero-KO mice (A) Sampling schedule after irradiation. (B) Accumulation of the irradiation-induced ssDNA as a DNA damage marker (magenta) in neurons in the dentate gyrus of the wild-type and the *Rp58* hetero-KO mice. *p < 0.05, Dunnett’s test vs the wild-type mice at 1hours after irradiation; n = 3 for each group. (C) Irradiation-induced accumulation of gamma-H2AX protein (gray) as a DNA damage marker in neurons in the dentate gyrus of the wild-type and the *Rp58* hetero-KO mice. *p < 0.05, **p < 0.01, Dunnett’s test vs the wild-type mice at 1hours after irradiation; n = 3 for each group. (D) Microglial activation detected by labeling microglia with Iba1 and activation with CD68 in the dentate gyrus of the wild-type and the *Rp58* hetero-KO mice. **p < 0.01, Dunnett’s test vs the wild-type mice at 1hours after irradiation; n = 3 for each group.

### Chronic treatment with minocycline prevents early aging phenotypes of Rp58 hetero-KO

We aimed to propose an effective therapeutic strategy for human aging by improving the early onset of cognitive dysfunction in our new aging mouse model, which has a DNA repair defect due to a human-like RP58 decline. Thus, the involvement of RP58 in DNA repair functions encouraged us to investigate the effect of minocycline, which has neuroprotective [15] and anti-inflammatory effects [14], on the early aging phenotypes of the *Rp58* hetero-KO mice. The *Rp58* hetero-KO and the control mice were administered minocycline for 1 month to 4–5 months in their drinking water (Figure 4A). The chronic minocycline treatment prevented the increase of ssDNA-positive and/or gamma-H2AX-positive neurons in the dentate gyrus of the *Rp58* hetero-KO mice at 4–5 months (Figure 4B, 4C). Furthermore, we examined the effect of minocycline on the early activation of microglia in the *Rp58* hetero-KO mice. Chronic treatment with minocycline prevented abnormal activation of microglia in the dentate gyrus of the *Rp58* hetero-KO mice (Figure 4D). Our findings indicate that minocycline is useful to prevent a series of age-related phenomena due to underexpression of RP58 in the dentate gyrus. As it is important to assess the effects of any treatment strategy on behavior, we also examined the effect of minocycline on the impairment of object location memory at 4–5 months in the *Rp58* hetero-KO mice (Figure 4E). Treatment with minocycline did indeed prevent this cognitive dysfunction observed in the *Rp58* hetero-KO mice but does not affect in the wild-type mice.

**Fig. 4.**
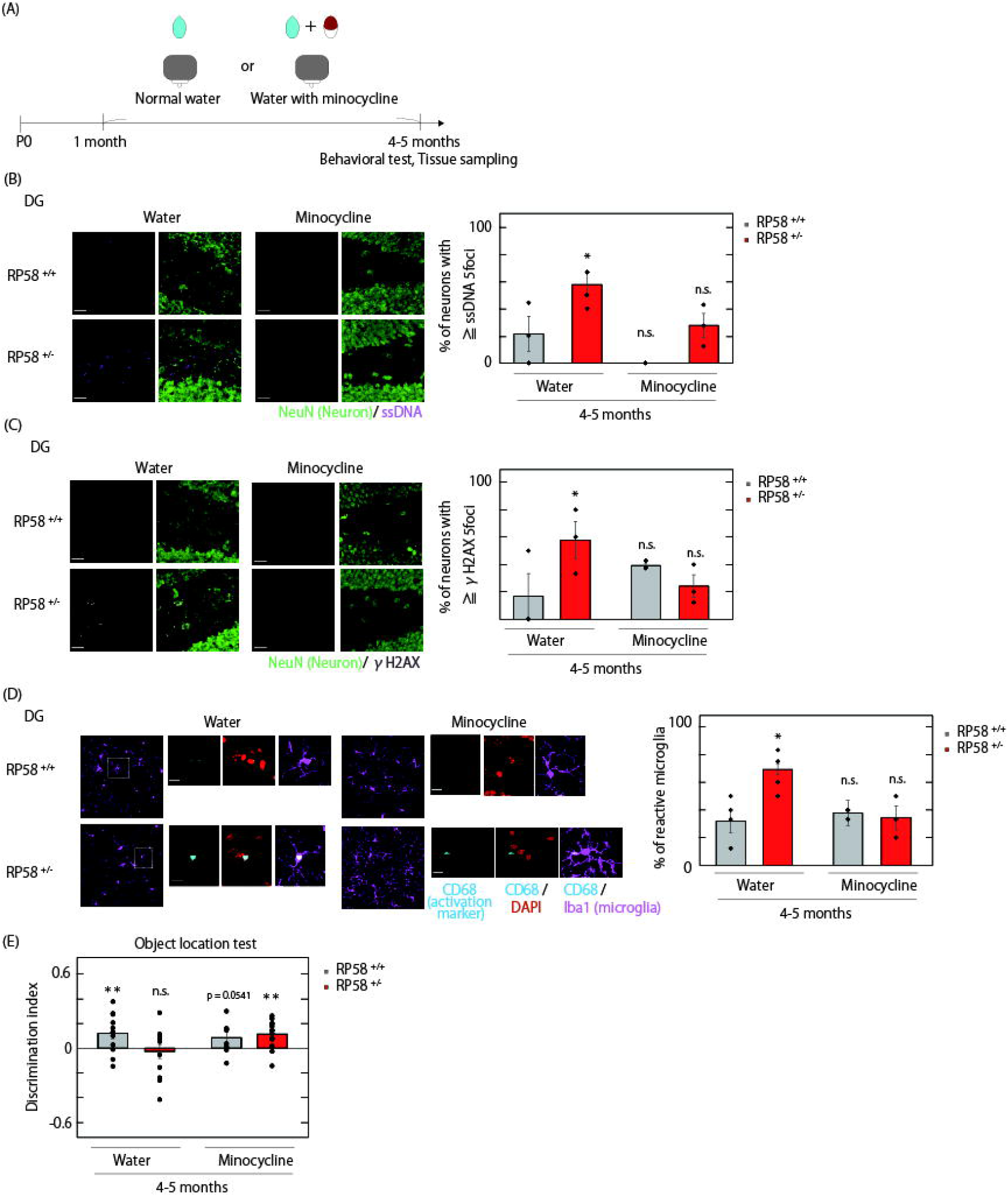
Minocycline treatment for human-like RP58 decline in mice (A) Experimental schedule of drug intervention. (B) The effect of minocycline on ssDNA accumulation (a DNA damage marker) in neurons in the dentate gyrus of the *Rp58* hetero-KO mice. *p < 0.05, Dunnett’s test vs the wild-type with normal water; n = 4 for both genotypes with normal water, and n = 3 for both genotypes with minocycline. (C) The effect of minocycline on gamma-H2AX protein accumulation (a DNA damage marker) in neurons in the dentate gyrus of the *Rp58* hetero-KO mice. *p < 0.05, Dunnett’s test vs the wild-type with normal water; n = 3 for each group. (D) The effect of minocycline on microglial activation in the dentate gyrus of the *Rp58* hetero-KO mice. *p < 0.05, Dunnett’s test vs the wild-type with normal water; n = 4 for both genotypes with normal water, and n = 3 for both genotypes with minocycline. (E) The effect of minocycline on the impairment of spatial memory in the *Rp58* hetero-KO mice. **p < 0.01, One sample t-test vs zero in the discrimination index: n = 12 for both genotypes with normal water, n = 10 for wild-type mice with minocycline at 4–5 months, and n = 16 for the *Rp58* hetero-KO mice with minocycline at 4–5 months. In this figure, we reposted the same data used in figure 1 and 2 as the control conditions treated with normal water according to the 3R rule.

These results consolidate that *Rp58* is involved in inflammation and DNA damage repair, and that its knockout leads to impairment of cognitive function. These findings suggest the availability of the *Rp58* hetero-KO mice as a novel mouse model of human-like early aging and provide a therapeutic strategy to prevent age-related phenomena by using minocycline.

## Discussion

Aging represents age-related decline in physiology. To extend the healthy lifespan and maintain quality of life, therapeutic strategies for age-related decline need to be developed. In this study, we generated a novel mouse model of human-like early aging phenotypes by hetero-KO of the *Rp58* gene and proposed a potential therapeutic strategy for age-related decline. Our *Rp58* hetero-KO mice showed early aging phenomena, including impairments in spatial cognition and DNA repair defects. In the electrophysiological analysis, the mutant mice exhibited an early aging phenotype related to the learning of object locations: the usual enhancement of theta power during the learning phase attenuated in the hippocampal CA1 region of the *Rp58* hetero-KO mice even at the adult stage. It should be noted that the *Rp58* hetero-KO mice only demonstrate some specific aspects of aging rather than complete age-related functional decline. This is because no significant effects of the *Rp58* hetero-KO were observed in other behavioral tests, such as the open-field test, the rotarod test, and the fear-conditioning test (Supplemental figure 2). The *Rp58* hetero-KO mice therefore represent a novel mouse model of human-like early aging specific to the processes underlying cognitive function. This model will be useful for the development of strategies to prevent mild cognitive impairment, which poses a major social challenge for aging society.

Our findings indicate that the cognitive dysfunction observed in the *Rp58* hetero-KO mice is due to impairments in DNA repair. Accumulation of DNA damage in the mutant mice was prevented by pretreatment with minocycline. Considering that autophagy induction is one of the key points of the application of minocycline [15], RP58 protein may play a role in autophagy in the DNA repair response. A decrease in autophagic activity is considered an age-related phenomenon and one of the causes of DNA damage accumulation [31]. RP58 protein is a transcription factor that represses the transcription of a series of genes which is reported to attenuate autophagy and apoptosis [32]. It has also been reported that autophagy is required for the survival and maturation of adult-born dentate gyrus neurons [33]. This evidence raises the possibility that autophagy dysfunction leads to DNA damage accumulation in the dentate gyrus neurons in the *Rp58* hetero-KO mice and emphasizes the importance of our current research on RP58 for our understanding of the aging mechanisms that are related to DNA damage accumulation.

Regarding another point of application of minocycline, it has been reported that minocycline selectively inhibits the activation of injurious microglia [14]. Microglia play a role in immune function, and microglial dysfunction is one of the typical age-related changes. With age, microglia change from a ramified morphological state to a de-ramified, spheroid formation with abnormal processes [34]. In the present study, microglial activation was detected in the wild-type mice at 12–18 months. In contrast, in the *Rp58* hetero-KO mice, microglial activation was already detected at 4–5 months.

Although microglia act protectively through inflammatory responses and phagocytosis at an early stage, they have an injurious effect at the late stage. In the present study, minocycline prevented the early activation of injurious microglia in the *Rp58* hetero-KO mice. Considering this point of application of minocycline and the effect of injurious microglia on neurons [35], microglia may play a crucial role in DNA damage accumulation in the neurons of the *Rp58* hetero-KO mice. It has been reported that dysfunction of mitochondria is an abnormal phenomenon that stems from DNA damage accumulation, induced by a metabolic imbalance, oxidative stress, inflammation, cellular injury, or cell death in many pathologies [36, 37]. Notably, abnormalities in mitochondria have been shown to be age-related [28]. In the *Rp58* hetero-KO mice, mitochondrial dysfunction in neurons may be caused by an abnormality in the immune and DNA repair system. Therefore, the RP58 deficit may induce impairments in autophagy, resulting in the accumulation of abnormal mitochondria and toxic proteins [38].

In summary, as a step from bench to bedside, we explored therapeutic strategies for aging by generating a unique mouse model. Interestingly, pretreatment with minocycline prevented DNA damage accumulation, abnormal activation of microglia, and impairment of spatial memory in our *Rp58* hetero-KO mice. Minocycline has not only direct effects on DNA damage and microglia but also indirect effects due to the relationship between DNA damage and inflammation [25, 39]. Minocycline has already been used against inflammatory diseases and, clinically, drug repositioning is easier than drug development. Our findings may thus be useful in developing strategies for preventing the decline in cognitive functions during aging.

In the present study, we established that RP58 plays a role in age-related DNA damage accumulation and dysfunction of the immune system. We advocated for the importance of DNA repair pathway and the immune system through RP58 in the underlying processes.

## Supporting information

Key resource table

## Acknowledgments

This work was partially supported by JSPS KAKENHI (grant number 18H02537) awarded to Haruo Okado, JSPS KAKENHI (17K16408), and by a Research Grant for Public Health Science and Suzuken Memorial Foundation awarded to Tomoko Tanaka, and by the Precursory Research for Innovative Medical Care (PRIME) from the Japan Agency for Medical Research and Development (AMED; 17937210) and AMED under Grant Number JP22gm4010019 awarded to Hikari Yoshitane.

The authors thank Dr. Nobuhiro Kurabayashi and Dr. Makoto Hashimoto for advice on the work presented here, and Kazuki Shiotani and Yuta Tanisumi for their help with the analysis of the electrophysiology data. We would also like to thank Editage (www.editage.com) for English language editing.

## Author contributions

T.T. and H.O. designed the research. H.Y and H.O supervised the study. T.T. performed and analyzed most experiments. K.E. performed the sample preparation for electron microscopy. Y.N. performed the complementary DNA microarray analysis of gene expression. H.M. helped with the setup for the electrophysiological experiment. S.H. and H.S. provided additional help with this study and made important suggestions during the process. T.T. generated all figures and tables. T.T. and H.Y. wrote the manuscript. H.O. edited the manuscript.

## STAR Methods

All experimental procedures were approved by the Animal Experimentation Ethics Committee of the Tokyo Metropolitan Institute of Medical Science (49040).

### Lead contact

Requests for information and resources not made available here should be directed to and will be fulfilled by the lead contacts, Tomoko Tanaka and Haruo Okado.

### Materials availability

This study did not generate new reagents.

### Experimental model and subject details

#### Animals

The original lines of RP58 mutant mice (the *Rp58* homo-KO mice) were established as a C57/BL6 congenic line (a genetic background strain) by repeated back-crosses, as described previously [9]. Because the *Rp58* homo-KO mice are embryonically lethal, the *Rp58* hetero-KO mice were chosen to investigate the process of aging. All mice were maintained under a 12:12 h light/dark cycle (lights on at 8:00 AM). All efforts were made to minimize the number of animals used and their suffering. All experiments were performed during adolescence (postnatal day 45 (P45)–P60), adulthood (P90–P120), or old age (P365–P545).

### Method details

#### Behavioral assessment

Male mice were allowed to habituate to the behavioral room for at least 1 hour before commencement of the behavioral test. Mice showing outlier behavior, i.e., >2 standard deviations or <2 standard deviations from the mean, were not analyzed as part of the behavioral test.

The object location test consisted of three phases as described previously [40]. All phases were performed under a light intensity of 10 lux. On day 1, the animals were placed in an empty square box (50 × 50 cm^2^) for 10 minutes. On day 2, the animals were placed in the same box for 10 minutes with two identical unfamiliar objects. On day 3, the animals were placed in the same box, with one of the two items being displaced to a novel location in the area. The discrimination index was calculated as the ratio of the time spent exploring the novel place to that spent exploring the familiar place: discrimination index = (novel place exploration time – familiar place exploration time)/(novel place exploration time + familiar place exploration time). On each day, the exploration time in each place was calculated automatically using a DVTrack video tracking system (Muromachi, Tokyo, Japan).

For the open field test, each mouse was placed in center of the apparatus (50 × 50 cm^2^) and was allowed to move freely for 30 minutes. Horizontal activity were collected every 10 minutes using a DVTrack video tracking system (Muromachi, Tokyo, Japan).

The rotarod test consisted of five trials with 1-hour intervals between trials. Mice were placed on a drum (3 cm diameter; MK-660B; Muromachi Kikai Co., Ltd., Tokyo, Japan) rotating at 1 and 4 rpm for 5 minutes to habituate to the rotarod prior to test. An ability of motor learning was evaluated in an accelerated rotarod test from 1 to 40 rpm over a 5-min period.

The fear conditioning test consisted of three phases as described previously [40]. On day 1, the animals were placed in a triangular box for 5 minutes under a light intensity of 30 lux. On day 2, the animals were placed into a square box with a stainless-steel grid floor and were allowed to explore the box freely for 2 minutes. Subsequently, a tone, which served as the conditioned stimulus (CS), was presented for 30 sec. During the last 1 sec of CS presentation, a 0.75 mA electric shock was applied, which served as the un-conditioning stimulus (US). Two more CS-US parings were presented with a 1-minute inter-stimulus interval under a light intensity of 100 lux. On day 3, the animals were placed into the same square box as used on day2 for 5 minutes under a light intensity of 100 lux. In each test, the percentage of freezing time and distance travelled (cm) were calculated automatically using ImageFZ software (O’Hara & Co., Tokyo, Japan). Freezing was taken to indicate that the mice remembered the box (context) in which they were exposed to the electrical shock.

#### Electrophysiology

Mice were anesthetized with isoflurane (1.5%) and implanted with a custom-designed electrode comprising two tetrodes in the hippocampal CA1 (1.8 mm anterior to bregma, 1.4 mm lateral to the midline, and 1.2 mm from the brain surface). Individual tetrodes consisted of four twisted polyimide-coated tungsten wires (single wire diameter 12.7 μm; California Fine Wire, Grover Beach, CA). One additional screw was threaded into the bone above the cerebellum for reference and grounding. The electrodes were connected to an electrode interface board (EIB-18, Neuralynx, MT) on a microdrive. The behavioral assessment was performed at least 1 week after surgery.

Electrical signals were obtained with open-source hardware (Open Ephys, Cambridge, MA) during the object location test for local field potential (LFP) recording at a sampling rate of 30 kHz. For synchronization with behavioral data, a transistor-transistor logic pulse was managed. After recording, electrode lesions were induced using 20 μA direct current stimulation for 10 seconds using one of the four tetrode leads.

#### Analysis of electrophysiology data

All data analyses were performed using the built-in software in MATLAB (MathWorks, Inc., MA). LFPs were down-sampled to 1,000 Hz and the data were extracted 1 second before and after contact with the objects. Theta power was calculated by taking the mean value of the power spectrum between 4 and 8 Hz. This value was normalized by the theta power of mice while walking.

#### Immunohistochemistry

Male and female mice in adolescence, adulthood, or old age were anesthetized with isoflurane and then sequentially perfused through the heart with PBS (Phosphate-buffered saline) followed by 4% paraformaldehyde (PFA) in PBS. Their brains were post-fixed, cryoprotected in 20% sucrose overnight and 30% sucrose over 2 days at 4□, and then embedded in optimal cutting temperature compound and frozen. Coronal sections of mouse brains were then cut using a cryostat (20 μm-thick).

Free-floating sections containing the hippocampus were activated by HistoVT One (Nakalai, Kyoto, Japan) at 70□ for 20 minutes, incubated in 0.4% Block Ase (DS Pharma Biomedical, Osaka, Japan) in PBS at room temperature (RT) for 20 minutes, and then incubated with primary antibodies (Supplemental Table 1). All primary antibodies were diluted 1:500 in PBS containing 0.3% Triton X-100. The sections were then incubated with secondary antibodies (Jackson, ME) (diluted 1:500 in PBS containing 0.3% Triton X-100) for 2 hours at RT. All section types were incubated with DAPI (Nacalai Tesque, Kyoto, Japan) for nuclear staining. Fluorescence micrographs of the sections subjected to immunostaining procedures were captured and digitized using a FluoView® FV3000 confocal laser scanning microscope (Olympus, Tokyo, Japan).

#### Analysis of immunostaining images

To quantify single-stranded DNA (ssDNA) and gamma H2AX in the cells, the number of cells with five or more nuclear ssDNA foci or gamma H2AX foci was counted [41]. For activated microglia analysis, CD68 antibody staining was performed. CD68 is a lysosomal-associated protein in macrophages/microglia and is associated with phagocytic cells. The Iba1 antibody was used as a marker for both resting and activated microglia. Microglia with aggregated CD68 expression were categorized as reactive microglia, and the percentage of reactive microglia was compared between groups [42, 43].

#### X-ray

Animals were irradiated with a dose of 0.3 Gray. At 1 hour, 6 hours, or 24 hours after irradiation, mice were anesthetized with isoflurane and then sequentially perfused through the heart with PBS followed by 4% PFA in PBS. Subsequent processes were similar to those described in the immunohistochemistry section.

#### Electron microscopy

Tissue preparation: In adulthood, male and female mice were anesthetized with isoflurane and then sequentially perfused through the heart with saline followed by 2% PFA and 2.5% glutaraldehyde in 0.1 mol/L PB (pH 7.4). Their brains were post-fixed, and coronal sections of the mouse brain containing the dentate gyrus and prefrontal cortex were then cut using a VT1200S microslicer (300 μm) (Leica, Wetzlar, Germany). Micro-dissected areas were post-fixed at 4□ for 2 hours in 2% osmium tetroxide in 0.1 mol/L cacodylate buffer. Tissue blocks were dehydrated using 50% ethanol for 5 minutes, 70% ethanol for 15 minutes, 80% ethanol for 15 minutes, 90% ethanol for 15 minutes, 100% ethanol at RT for 20 minutes × 3, and were subsequently infiltrated with propylene oxide for 10 minutes × 3, with 1:1 solution of propylene oxide: epoxy resin for 2 hours and epoxy resin overnight. Each block was placed flat on glass microscope slides and horizontally mounted on gelatin capsules (Lilly, IN). After embedding, each block was polymerized in epoxy resin (EPON 812, TAAB, Berkshire, England) at 60 °C for 48 hours. Polymerized blocks were trimmed to a section containing the dentate gyrus and prefrontal cortex using an ultramicrotome PowerTomeX (RMC Boeckeler, AZ). Semi-thin (1 μm) sections were stained with toluidine blue and used to guide further cutting of the specimen block to ultra-thin sections (50–80 nm). Ultra-thin sections were placed in formvar-coated single-slot grids. After staining with uranyl acetate and lead citrate, ultra-thin sections were observed under a JEM-1400 transmission electron microscope (JEOL, Tokyo, Japan) equipped with a bottom-mounted charge-coupled device camera and subsequently processed using ImageJ.

Quantification of ultrastructural defects: Quantification of mitochondrial ultrastructural defects was performed from transmission electron microscopy images at a magnification of 10,000 ×. Although mitochondria are sparse within astrocytic processes of the dentate gyrus, the criteria used by Siskova et al.^43^ [44] for the identification of astrocytes and neuronal cells were applied to ensure that predominantly, if not exclusively, neuronal mitochondria were included. Every mitochondrion of neuronal origin was analyzed and, based on its ultrastructural appearance, classified into one of the following categories according to Siskova et al. ^37^[37]: intact mitochondria with normal-appearing cristae (Type I); abnormal mitochondria with either swollen, irregular, or whirling cristae (Type II); mitochondria with a discontinuous outer membrane or deficient cristae (Type III); and mitochondria with both swollen and deficient cristae or both a discontinuous outer membrane and swollen cristae (Type IV). At least 14 mitochondria from each region were analyzed for each animal.

#### Drug treatment

Minocycline was administered to the animals’ drinking water from weaning to the day before the behavioral test or perfusion in the wild-type and the *Rp58* hetero-KO mice. Minocycline was dissolved in the filtrate water at 0.015 mg/mL.

#### Transcriptome analysis

Three independent total RNA samples from each group were mixed and purified using a RNeasy Plus Universal Kit (Qiagen, Hilden, Germany). RNA quality was assessed using a 2100 Bioanalyzer (Agilent Technologies, Santa Clara, CA). Cy3-labeled complementary RNA was prepared using a Low Input Quick Amp Labeling Kit (Agilent Technologies) in accordance with the manufacturer’s protocol. Samples were hybridized to the SurePrint G3 Mouse Gene Expression v2 Microarray (G4852B; Agilent Technologies). Thereafter, the array was washed and scanned using the SureScan Microarray Scanner (Agilent Technologies). Microarray images were analyzed using Agilent’s Feature Extraction software (Agilent Technologies) with default settings for all parameters. Data from each microarray analysis were normalized by shifting to the 75th percentile without baseline transformation.

#### Statistical data analysis

Group data are presented as the mean ± SEM. The statistical significance of between-group differences was assessed by One sample t-test, Tukey–Kramer test, Dunnett’s test, or paired t-test, using JMP software (SUS Institute, NC).

## Supplemental material

**Supplemental fig. 1** (A) Venn diagrams depicting an overlap in hippocampus genes exhibiting a < 0.5-fold or >1.5-fold expression change compared with the wild-type mice at 4–5 months. n = 3 for each group. (B) Scatter plot depicting 103 up-regulated genes. (C) Top 4 pathway enrichment analysis of commonly expressed genes.

**Supplemental fig. 2** Analysis of other behavior in the *Rp58* hetero-KO mice (A) Horizontal activity was calculated using an open-field test in the wild-type and the *Rp58* hetero-KO mice at 4–5 months. Horizontal activity was calculated per 10 minutes. Two-way repeated ANOVA followed by Holm’s test; n = 14 for the wild-type mice at 4–5 months, n = 11 for the *Rp58* hetero-KO mice at 4–5 months. (B) Motor learning test of the wild-type and the *Rp58* hetero-KO mice at 4–5 months. The latency to fall was recorded for 5 consecutive days. Two-way repeated ANOVA followed by Holm’s test; n = 12 for the wild-type mice at 4–5 months, n = 11 for the *Rp58* hetero-KO mice at 4–5 months. (C) Contextual fear memory test of the wild-type and the *Rp58* hetero-KO mice at 4–5 months. Freezing time was calculated per 1 minute. Two-way repeated ANOVA followed by Holm’s test; n = 23 for the wild-type mice at 4–5 months, n = 18 for the *Rp58* hetero-KO mice at 4–5 months.

