## Supplementary figures and images for "A tetracycline antibiotic minocycline prevents early aging phenotypes in mice heterozygous for RP58"

### Supplimental figure 1

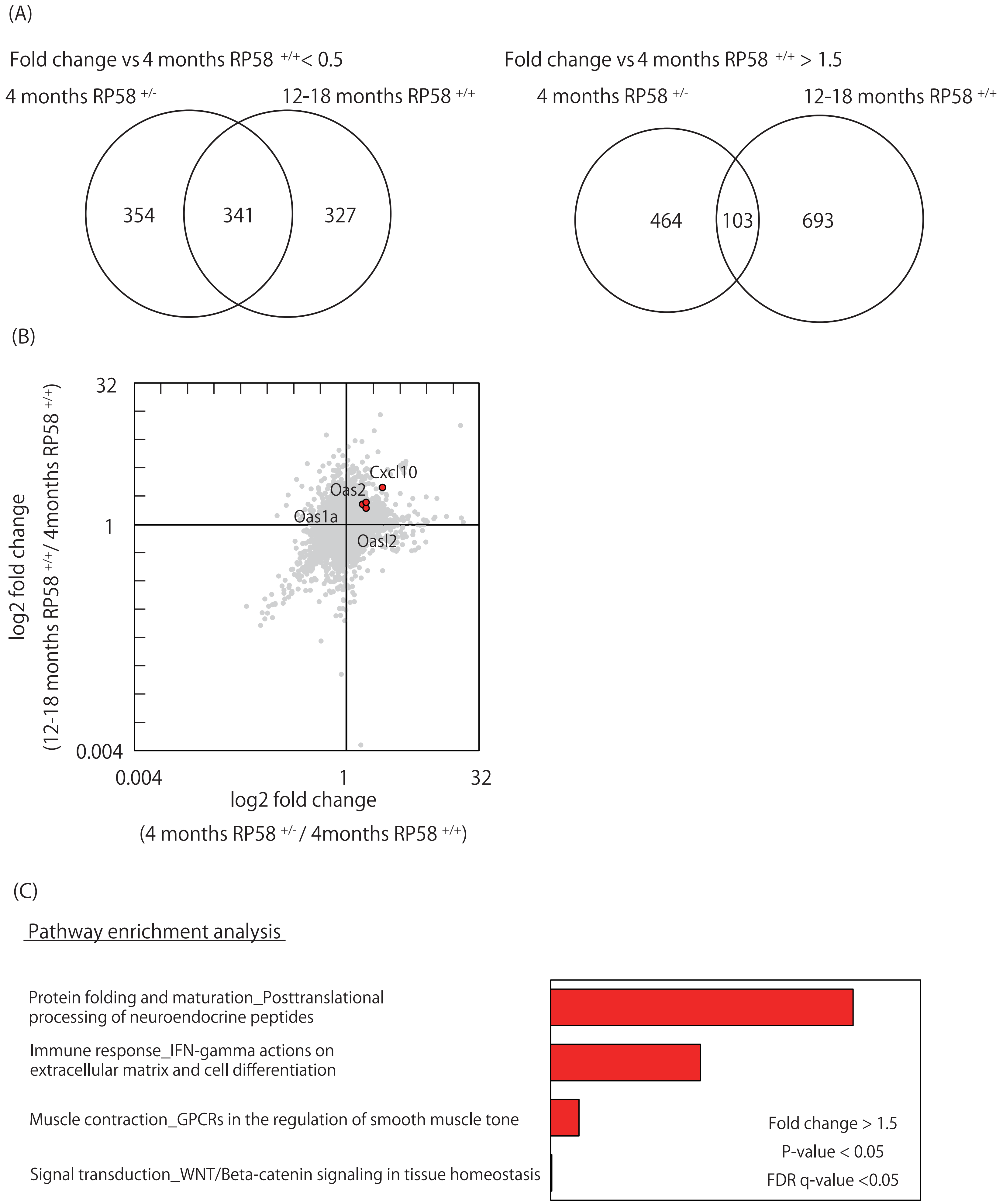

### Supplimental figure 2

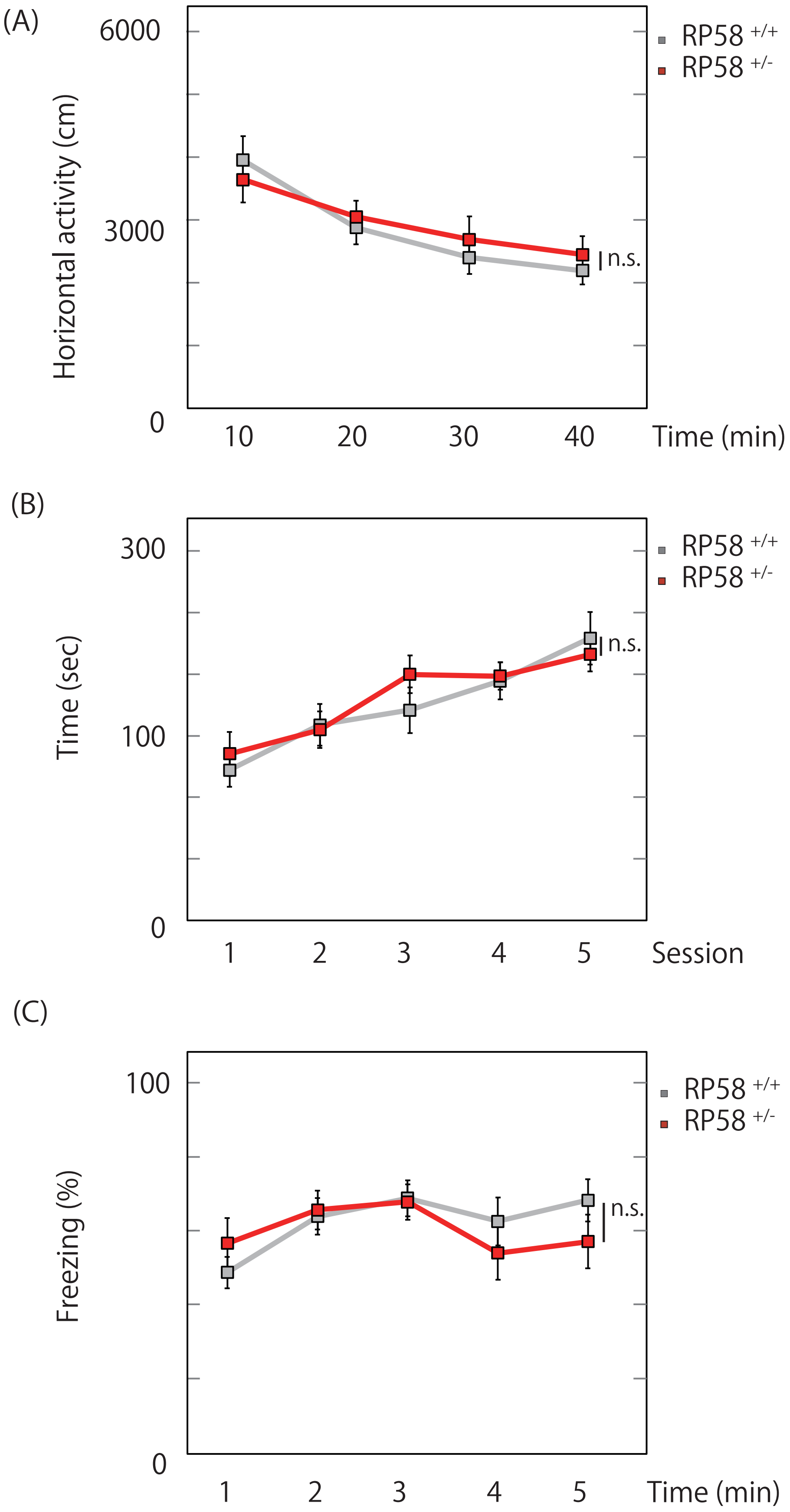
